# Hydrogen Peroxide Levels in Freshly Brewed Coffee and the Effects of Storage

**DOI:** 10.1101/2020.08.08.242552

**Authors:** Sannihith N. Uppu, Bianca K. London

## Abstract

One of the world’s most consumed beverages, coffee has its origins as early as the 15^th^ century Ethiopia. Although there are studies on caffeine and other components of coffee such as cafestol and kahweol, up until recently knowledge of the presence of hydrogen peroxide (H_2_O_2_) in coffee was confined to the scientific community and some informed public. It is a general belief that H_2_O_2_ is formed only after long periods of storage or with certain roasting practices. The present study is focused on dispelling the myths of H_2_O_2_ in coffee. We first measured H_2_O_2_ in freshly brewed coffee from different companies using the ferrous oxidation-xylenol orange binding (FOX) assay. Following this, we examined the time-dependent accumulation of H_2_O_2_ and its changes with temperature. Further, H_2_O_2_ was estimated in coffee obtained from several local vendors. Contrary to the general belief that the accumulation of H_2_O_2_ is an aging phenomenon of coffee, we found this toxicant even in freshly brewed coffee. This was true for all brands tested, and the H_2_O_2_ content increased upon storage. The highest increase was seen in coffee stored on the hot plate compared to the ones kept at room temperature (22-25 °C) or in the cold (0-4 °C). The H_2_O_2_ content of coffee from different vendors ranged between 0.29 and 0.82 mM, which is 5- to 20-fold higher than the typical H_2_O_2_ concentrations at which significant cytotoxic effects have been reported for assay systems using the human Fanconi deficient (PD20 FANCD2−/−) fibroblasts and other cell types. Our findings are deemed to shine new light on the probable toxic effects of a commonly consumed beverage like coffee, and the time and temperature dependent variations of keeping. While there are documented benefits of consumption of coffee, the possible H_2_O_2_-medicated toxic effects are critical and should be considered. Future studies are warranted to delineate the contribution of H_2_O_2_ in the healthy wellbeing of individuals who consume coffee extensively.

## Introduction

Coffee is one of the most consumed beverages in the world. An estimated 50% of all adults consume it daily. Coffee is produced from ground coffee plant beans and contains abundant amounts of polyphenolic compounds. Consumption of this polyphenol rich beverage is believed to have beneficial health effects on the human body but may also have adverse effects due to the production of hydrogen peroxide, H_2_O_2_ (1-5). Experimental studies with human volunteers have shown increased H_2_O_2_ levels in urine (6-8). These increased levels of H_2_O_2_ are attributed to the autooxidation of polyphenols in the bladder and do not come from coffee as such. The H_2_O_2_ in coffee accounts for many of the observed cytotoxic, genotoxic and mutagenic effects of coffee in isolated cell culture systems (9-14).

In this study, we show that significant amounts of H_2_O_2_ are present in coffee from different vendors. Depending on the temperature at which coffee is stored, the H_2_O_2_ concentration in coffee reaches as high as 1 mM, which is 5- to 20-fold higher than the typical H_2_O_2_ that induces signaling including apoptosis in most mammalian cell types in culture (4, 7, 10). The high concentrations of H_2_O_2_, particularly in stored coffee, begs the question of what unforeseen health impacts drinking large quantities of coffee regularly could have.

## Materials and Methods

### Chemicals and reagents

Xylenol orange and H_2_O_2_ (30%) were purchased from Sigma-Aldrich (St. Louis, MO). Ferrous ammonium sulfate and all other chemicals used in the study were of analytical grade. Water used in the preparation of reagents and brewing of coffee was deionized to a final resistance of 18.0 mΩ/cm or higher.

Hydrogen peroxide concentration in the stock solution was determined spectrophotometrically using €_240 nm_ of 43.6 M^-1^cm^-1^ (15). The working standards of H_2_O_2_ (0-500 µM) were prepared by serial dilution using water.

FOX reagent was prepared as described by Gay and Gebicki (16) with some minor modifications. Briefly, 19 mg of xylenol orange and 78 mg of ferrous ammonium sulfate were dissolved successively in 20 mL of 0.2 M perchloric acid. The volume of the mixture was then made up 100 mL with water.

### Brewing of Coffee

Ground coffee (10 g) was brewed using 350 mL (*ca*. 11 Oz) of deionized water. The brewing typically took 5 min to complete. A small portion (2-5 mL) of the brewed coffee was collected in a test tube and cooled under running tap water for 1min.

### Determination of H_2_O_2_

Aliquots (20 to 50 µL each) of cooled coffee was added to 950 µL of FOX reagent. The volume was brought to 1 mL by addition of required amounts of water (typically 0 to 30 µL), mixed well on a vortex for 10-15 s, and incubated for 5 min at the room temperature (22-25 °C). At the end of the incubation period, the absorbance of the coffee-FOX reagent mixtures was measured at 560 nm using a Genesis 10S UV-Vis spectrophotometer. Reagent blanks were set out by mixing 50 µL of water with 950 µL of the FOX reagent and measured after 5 min of incubation at the room temperature.

A standard curve for the H_2_O_2_ determination involved mixing 50 µL of 100 to 500 µM H_2_O_2_ with 950 µL of the FOX reagent as described above (final concentration H_2_O_2_: 5-25 µM). Reagent blanks were set out as described above. The absorbance of the blank at 560 nm was subtracted from the respective H_2_O_2_ standards before the construction of the standard curve.

### Effects of Storage

Freshly brewed coffee after cooling under the running water (see above) was stored on ice/water mixtures or left at the room temperature for periods up to 3 h. In some cases, the coffee was left on the hot plate (*ca*. 70 °C) of the coffee machine and analyzed for the presence of H_2_O_2_ at various periods of storage (15 min to 3 h) as described above.

All H_2_O_2_ determinations were performed in either duplicate or triplicate and averaged the individual readings.

## Results

Table I shows the typical concentrations of H_2_O_2_ in freshly brewed coffee samples from three different vendors. The concentrations are typically in the range of 0.14 to 0.17 mM. The brand and the type roasting the coffee beans underwent at the manufacturing facility did not seem to influence the concentration of H_2_O_2_ in the freshly brewed coffee.

**Table I.** H_2_O_2_ content in freshly brewed coffee from three different vendors

| Brand | H<sub>2</sub>O<sub>2</sub> (mM) |  |
| --- | --- | --- |
|  | Sample 1 | Sample 2 |
| Folgers (classic roast, medium) | 0.16 | 0.17 |
| Great Value (classic roast, medium) | 0.14 | 0.15 |
| Community (breakfast blend, medium-dark) | 0.17 | 0.17 |

Note: H<sub>2</sub>O<sub>2</sub> content analyzed by the FOX assay as outlined in the Methods section.

For a given brand of coffee, the concentration of H_2_O_2_ increased upon storage (Table II). The buildup of H_2_O_2_ with time was dependent on the temperature at which the coffee was stored. When left on the hotplate provided with the coffee machine, the extent of increase over a period of 3 h was found to be as high as 0.55 mM, about 200% more than the starting concentration of H_2_O_2_. For the same period, the increase in the H_2_O_2_ concentration was only about 100% more (0.35 mM) if the coffee was stored at typical household temperatures of 23-25 °C. The best storage which resulted in a minimal increase of 50% or less in 3 h period appeared to require refrigerated temperatures of 32-41°F.

**Table II.**
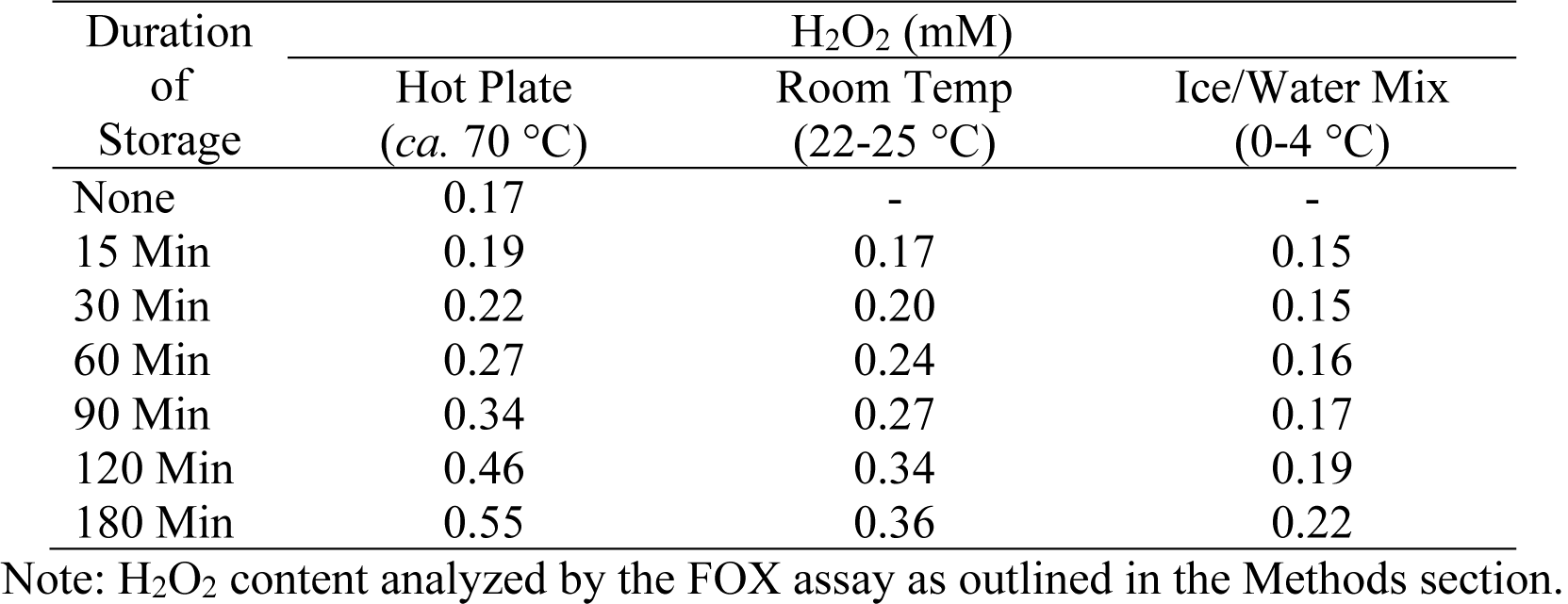
Time-dependent increase in H_2_O_2_ concentration in freshly brewed coffee.

| Duration of Storage | H<sub>2</sub>O<sub>2</sub> (mM) |  |  |
| --- | --- | --- | --- |
|  | Hot Plate (*ca.* 70 °C) | Room Temp (22-25 °C) | Ice/Water Mix (0-4 °C) |
| None | 0.17 | - | - |
| 15 Min | 0.19 | 0.17 | 0.15 |
| 30 Min | 0.22 | 0.20 | 0.15 |
| 60 Min | 0.27 | 0.24 | 0.16 |
| 90 Min | 0.34 | 0.27 | 0.17 |
| 120 Min | 0.46 | 0.34 | 0.19 |
| 180 Min | 0.55 | 0.36 | 0.22 |

Note: H<sub>2</sub>O<sub>2</sub> content analyzed by the FOX assay as outlined in the Methods section.

Figure 1 shows data on brewed coffee samples collected from different vendors. The concentrations ranged from 0.29 to 0.82 mM. Although we tried to analyze coffee samples in less than 1 h after they were collected from the respective vendors, we suspect that were lot of variables such as the process of brewing and how long the coffee was stored before it was brought? Thus, while we are not prescriptive about the concentration of H_2_O_2_ in these samples, we believe this provides unequivocal evidence for the presence of H_2_O_2_ in coffee wherever it comes from.

**Fig. 1.**
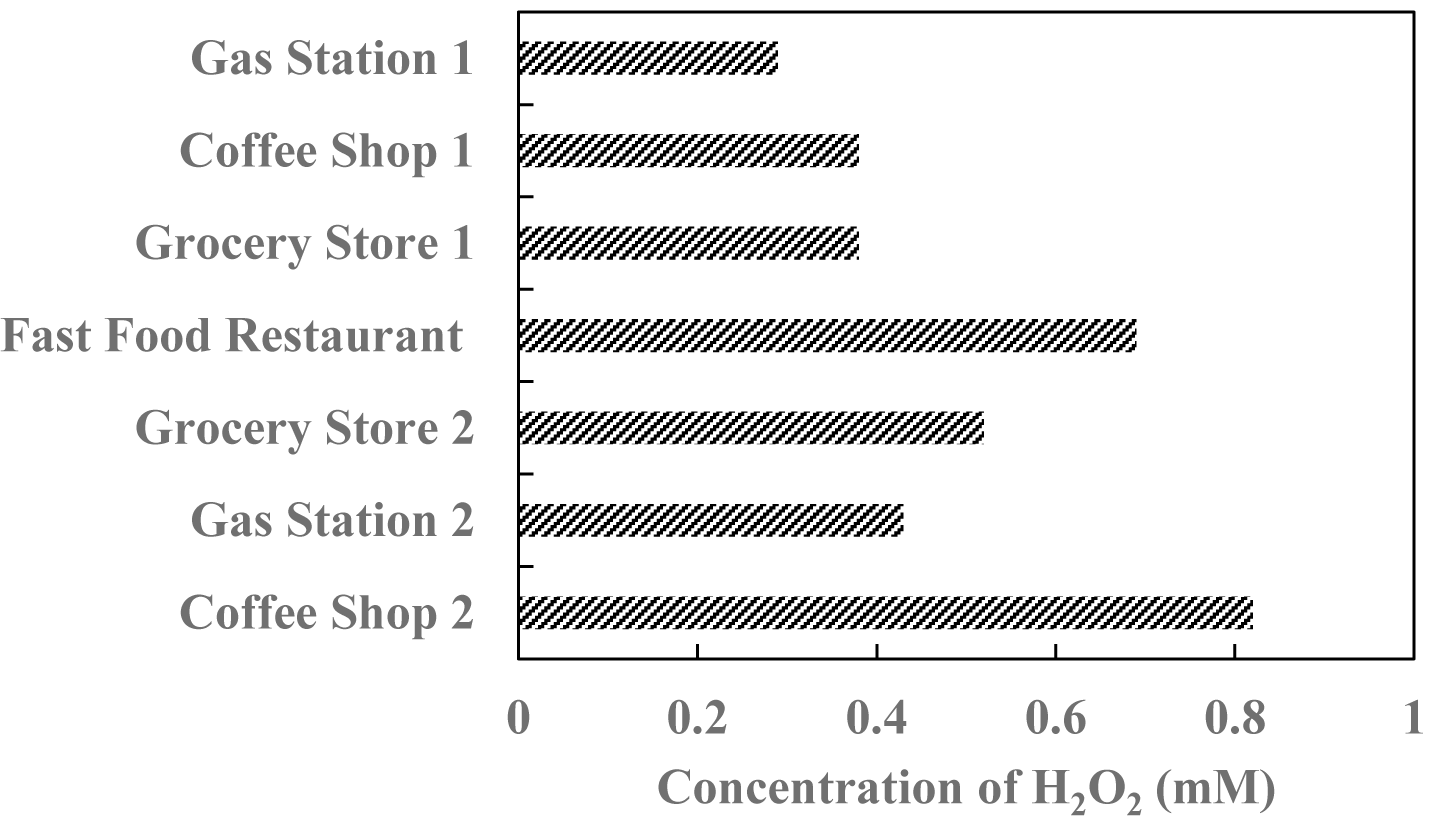
H_2_O_2_ content in coffee samples collected from different vendors. Each data point resents an average of two or three determinations.

## Discussion

The results of the present study reveal that significant amounts of H_2_O_2_ are present even in freshly brewed coffee. This is true for all three brands tested. Further, the H_2_O_2_ content of coffee increases upon storage. The highest increase is seen with the coffee stored on the hot plate compared to the ones at room temperature or in the cold. Though the coffee samples from different brews were relatively the same, the same cannot be said for the different vendors, which ranged from 0.29 mM to 0.82 mM. While the relative contribution is not known, we believe H_2_O_2_ in the coffees may be coming from the Millard reaction products formed during the roasting process (9, 12, 17) and/or redox cycling of hydroxyhydroquinone (8, 18) and polyphenolic constituents including chlorogenic acid (2, 3).

This brings forth the topic of the physiological significance of the H_2_O_2_ in coffee. We believe there is no simple answer to this question. Several cell cultures studies have demonstrated H_2_O_2_-mediated apoptosis or necrotic cell death at concentration of H_2_O_2_ of 50 µM or more (19) which are only a fraction (about one-fifth to one-twentieth) of the values reported here. H_2_O_2_ *per se* is not very reactive but it can result in the formation of hydroxyl radicals (HO^•^) through Fe(II)-mediated Fenton reactions or through non-classical Fenton reactions involving other transition metals like Cu(I). Though these reactions H_2_O_2_ could promote damage to DNA and host of other biomolecules (4, 5, 14, 19). However, as enumerated by Halliwell and his colleagues (19), there are a very efficient mechanisms for removal of H_2_O_2_ in the body. Therefore, it is very unlikely that drinking coffee *per se* may cause an increase in serum or tissue H_2_O_2_.

It is possible though the urinary H_2_O_2_ levels may be elevated following coffee consumption (6, 7). The increase, in part, has been thought to be due to the diffusion of H_2_O_2_ from coffee into oral cavity and the upper gastrointestinal tract (7). A major contributor to the urinary H_2_O_2_ after coffee drinking could be due to autooxidation of hydroxyhydroquinone in the bladder (8, 18, 19).

Drinking coffee on regular basis has been shown to have numerous benefits. These include prevention of cardiovascular disease, reduced risk of type-2 diabetes, and prevention of gall stone formation and gallbladder disease (20-23). Coffee shows antimicrobial activity against several species including *E. Coli, Listeria innocua*, and *Heliobacteria pylori* and this activity of coffee has been demonstrated mainly due to presence of H_2_O_2_ (13). Coffee consumption has been of some concern that it may increase the risk of developing bladder cancer in adults and leukemia in children of mothers who drank coffee during the pregnancy (24, 25).

Overall, the benefits of coffee outweigh the undesired affects. We are however particularly concerned about those who drink several cups of coffee in day and brew coffee over extended hours. This may add to the body’s oxidant stress.

## Acknowledgments

The authors acknowledge the support from the US Department of Education (US DoE; Title III, HBGI Part B grant number P031B040030). Its contents are solely the responsibility of authors and do not represent the official views of US DoE.

